# Use of plant growth-promoting bacteria to stimulate the growth and development of rice (*O*. *sativa* L.) and bean (*P*. *vulgaris* L.) cultivars

**DOI:** 10.1101/2025.10.18.683209

**Authors:** Luis Sierra, Domenico Pavone-Maniscalco, Oscar Valbuena

## Abstract

Current fertilization practices are unsustainable from both an environmental and a human health perspective. A promising solution has emerged in plant growth-promoting bacteria (PGPB), which can act hand in hand with synthetic fertilizers to reduce their environmental costs. This study seeks to isolate, characterize, and evaluate the ability of nitrogen-fixing bacteria (NFB) as a potential biofertilizer of important crops, such as bean and rice cultivars. 18 NFB strains were isolated from the rhizosphere of beans, lentils, and rice plants, with a few isolated from nodules or topsoil. To identify elite strains, primary growth-promoting mechanisms were determined, including mineral solubilization, N2-fixation ability, and production and secretion of indole-3-acetic acid (IAA). Of the 18 initial isolates, BN03, BN09, BN16 and BN18 showed the greatest potential as biofertilizers. These isolates were used individually and in consortium as bioinoculants for bean and rice plants in pot experiments, showing significant improvements in every biometric parameter measured. These results join others in showcasing the potential of beneficial microorganisms to promote the development and growth of important crops in the transition to sustainable agriculture.

## 1. INTRODUCTION

Food security worldwide depends on the use of synthetic fertilizers (Walling & Vaneeckhaute, 2020). For instance, about half of the global population is currently fed with crops grown with N-fertilizers (Erisman *et al*., 2008). Such a heavy use of synthetics is considered to be the catalyst of the explosive growth of human populations in recent times (Smil, 1999), allowing an increase from 1,6 billion in 1900 to more than 8 billion today (United Nations and Division, 2017). Although population growth has slowed down in recent decades, the Food and Agriculture Organization (FAO) still predicts that the world’s population will exceed 9 billion by the year 2050 (Conforti, 2011), forcing agricultural production to increase by 1,5% per year to cope with newer and bigger global food demands (Tilman *et al*., 2011; Walling & Vaneeckhaute, 2020).

Alongside the growth in population and agriculture, the production and use of synthetic fertilizers has also increased significantly (Bodirsky *et al*., 2014; FAO, 2018). The FAO reported an annual increase in nitrogen, phosphorus, and potassium fertilizer supply of 1%, 2,1%, and 3,8% between 2016 and 2022 (FAO, 2019), and further increases are expected in the near term. Although fertilizers play a pivotal role in the global food supply, they are also problematic (Gao & Cabrera, 2023). Their use has introduced a series of environmental and human health risks that make their current supply chain unsustainable in the foreseeable future (Zhang *et al*., 2015; Menegat *et al*., 2022), These adverse environmental and health impacts include greenhouse gas emissions (GHG), eutrophication, tropospheric pollution, soil salinization and alkalization, stratospheric ozone depletion, and reductions in soil biodiversity (Galloway, 2003; Davidson & Kanter, 2014; Ma *et al*., 2021; Jain, 2023). These impacts stem largely from the escape of reactive forms of N, P, and K from agricultural soils through leaching or volatilization (Zhang *et al*., 2015).

Global greenhouse gas emissions continue to rise at a time when they should be falling rapidly. The global surface temperature is now 1,15°C higher than pre-industrial levels (IPCC, 2023), with current atmospheric CO_2_ concentrations not previously seen in the past 14 Ma, according to recent paleo-CO_2_ estimates (CenCO2PIP, 2023). Agriculture is responsible for roughly 12% of all global net GHG emissions (Chataut *et al*., 2023), mainly composed of carbon dioxide (CO_2_) and nitrous oxide (N_2_O), along with notable emissions of ammonia (NH_3_) and nitrogen oxides (NO_x_). Although mitigation actions can reduce the emissions profile of chemical fertilizers (Gao & Cabrera, 2023), novel alternatives can be adopted to minimize the environmental pollution. The use of plant-beneficial microorganisms is a promising technology that is already available at the commercial level (Glick, 2012).

These plant growth-promoting bacteria (PGPB) have mechanisms that increase nutrient availability for plants, stimulate development by secretion of plant-growth regulators, increase the resistance to pathogens and pests, and improve abiotic resistance (Bhardwaj *et al*., 2014; de Souza *et al*., 2015; Poria *et al*., 2022). These benefits make the production and use of these biofertilizers an attractive tool that can be used in addition to synthetic fertilizers to increase crop yields, while paying a lower environmental and human-health cost (Singh *et al*., 2011; Zvinavashe *et al*., 2021). In response to climate change and the transition to sustainable agriculture, the market for biofertilizers is in constant search of new bacterial products that farmers can use to promote their crops (Fiodor *et al*. 2023). Therefore the present study was designed to isolate, characterize, and select high-performing nitrogen-fixing bacteria (NFB), a popular type of PBGB. The elite bacterial strains were then used in greenhouse assays to evaluate their potential to promote the growth and development of economically important crops, such as bean (*Phaseolus vulgaris* L.) and rice (*Oryza sativa* L.).

## 2. MATERIALS AND METHODS

### 2.1 Biological materials

Rhizosphere samples were collected the roots of wild bean (*P. vulgaris*), rice (*O. sativa*), lentil (*L. culinaris*), and common pea plants (Fabaceae sp., Poaceae sp.), located on different municipalities of Valencia, Carabobo state, Venezuela. The plants were uprooted and shaken to remove bulk soil, placed in sterile plastic bags, and stored at 4°C until they were processed (Venieraki *et al*. 2011). Nodule samples were obtained from bean trap hosts that derived from commercially sourced seeds. Topsoil samples were taken from the first 5 cm of the soil adjacent to the wild plants used for rhizosphere sampling. The soil samples were placed in sterile plastic bags, and stored at 4°C until processing (Ha & Chu, 2020). Bean and rice seeds used for pot experiments were donated from local agricultural vendors.

### 2.2 Reagents and growth media

All reagents used throughout the study were obtained from the Center for Applied Biotechnology (CAB), University of Carabobo, Venezuela. Rhizobacteria and rhizobia were cultured on Ashby and Yeast Mannitol Agar (YMA), respectively. Putative *Azospirillum* strains were cultured in semisolid N-free agar (NFb) and congo red agar (RC). Media composition can be found at Amaresan *et al*., (2022). N-free media was validated by two negative controls (*Escherichia coli* ATCC 8739, and *Staphylococcus aureus* ATCC 6538). Bidistilled water with a conductivity of < 0,1 µS/cm was used to minimize mineral contamination.

### 2.3 Isolation of nitrogen-fixing bacteria

#### 2.3.1 Rhizospheric and free-living bacteria

Ten grams of soil were collected aseptically and suspended in 90 mL of 0,85% NaCl solution. The mixture was stirred at 150 rpm for 30 min, and serial dilutions were diluted up to a final factor of 10^−6^. A 100 µL aliquot of each dilution was: (i) surface-streaked onto Ashby agar plates supplemented with sucrose and D-mannitol (Brenner *et al*. 2005), and (ii) inoculated in NFb tubes to screen for *Azospirillum* spp. Ashby plates and NFb tubes were incubated for one week at 28°C, and 37°C, respectively (Reis *et al*. 2015; Amaresan *et al*., 2022). Alkalized NFb tubes with a white diffuse pellicle below the surface were streaked onto RC plates (Cáceres, 1982). After three days of incubation at 37°C, small scarlet colonies with undulate edges were considered to be indicative of *Azospirrilum* spp. Isolation was carried out in aerobiosis for both types of NFB.

#### 2.3.2 Rhizobia

Bean seeds used to grow trap hosts were surface-sterilized with 70% ethanol for 30 s, and then with 3% sodium hypochlorite for 2 min. Finally, they were washed 2 times with sterile distilled water (Artigas *et al*. 2019). The seeds were pre-germinated in darkness for 48 hours on water-soaked, sterile Whatman grade filter paper. Ten pre-germinated seeds were sown in glass jars containing soil supplemented with commercial compost. The moisture level was maintained by adding sterile tap water. Plants were grown for 4 weeks under ambient conditions. Harvested nodules were surface-sterilized similarly as described before with seed sterilization (Artigas *et al*. 2021). Treated nodules were crushed in 500 µL of 0,85% of NaCl to obtain bacterial suspensions, which were serially diluted to 10^−6^. 100 µL aliquots of each dilution were pour-plated and streaked onto YMA, and plates were incubated under anaerobiosis at 28°C for one week using a GasPak system (Amaresan *et al*., 2022).

### 2.4 Characterization of NFB isolates

The biochemical profile and morphological traits were determined for every strain, along with their metabolic profile. The shape, Gram-type, sporulation, and cyst formation ability were determined in triplicate for all isolates, using 24 h fresh cultures (Beveridge *et al*. 2007). Gram-response was confirmed by L-aminopeptidase testing (Hernández *et al*. 1991). For biochemical testing, sugar fermentation assays (D-xylose, D-glucose, lactose, D-sorbitol, D-fructose, D-mannitol, and sucrose), motility tests, utilization of organic salts (citrate), and a enzymatic assay (catalase) were carried out, all in triplicate and using 24 h fresh cultures. All biochemical testing was performed according to the manual of MacFaddin (2003).

### 2.5 Screening for plant-growth promoting effects

#### 2.5.1 Quantification of nitrogen-fixing ability

The salycilate method was used to determine the total ammoniacal nitrogen (TAN) fixed by each isolate Giner-Sanz *et al*. (2021), which served as an indirect measure of nitrogenase activity. Fresh cultures were adjusted to 0,5 McFarland with a sterile NaCl solution. 1 mL of each suspension (10^8^ cells) was added to 9 mL of Ashby sucrose broth. Tubes were incubated for one week at 30°C. After incubation, the cell-free supernatant was recovered by centrifugation at 10000 *g* for 10 min. For each isolate, 5 mL of the supernatant was mixed with 600 µL of solution S1 (salycilate-nitroprussiate mixture) and 1 mL of solution S3 (alkaline hypoclorite mixture), after which samples were kept in darkness for 45 min. The absorbance at 640 nm was measured with a UV-Vis spectrophotometer (Fisher Scientific Genesys 10UV). Quantification of TAN was carried out with a calibration curve of NH_4_Cl (0-125 mg/mL). Each isolate was tested in triplicate.

#### 2.5.2 Bacterial mineral solubilization

The capacity of each bacterial strain to solubilize non-readily available forms of P, K and Zn was determined. Solubilization was qualitatively assessed with the plate assay method using media supplemented with minerals containing P, K, and Zn. P-solubilization was screened on NBRIP agar supplemented with Ca_3_(PO)_2_ (5 g/L) and 0,25% bromothymol blue to test the production of organic acids Nautiyal (1999). K-solubilization was assessed on Aleksandrow agar supplemented separately with feldspar and muscovite (3 g/L each), as described by (Pérez-Pérez *et al*., 2021). Zn-solubilization was screened in minimal media with added ZnO (2 g/L), as described by Yasmin *et al*. (2021). The experimental protocol was the same for each mineral. Fresh cultures were adjusted to 0,5 McFarland, of which 10 µL aliquots were spotted in their corresponding media, and incubated for one week at 30°C. The appearance of transparent halos around the bacterial colonies was indicative of positive activity. Each isolate was tested five times. The solubilization index (SI) was calculated for each nutrient according to Waday *et al*. (2021), using equation (1):

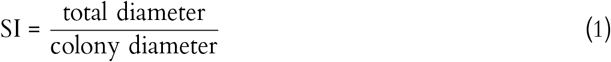

According to Teles *et al*. (2024), solubilization capacity is considered low if SI < 2, medium if 2 ≤ SI ≤ 4, and high if SI > 4. Additionally, the amount of solubilized P was quantified using the ascorbic acid method described by Murphy & Riley (1962). After incubation in NBRIP broth at 30°, the cell-free supernatant was recovered after centrifugation at 10000 g for 10 min. For each strain, 0,3 mL of supernatant was mixed with 0,7 mL of solution III (an acid mixture of ammonia molybdate with ascorbic acid). The mixtures were incubated in darkness for 20 minutes at 45°C, after which the absorbance was measured at 820 nm with a UV-Vis spectrophotometer. To quantify the amount of phosphate solubilized, a calibration curve of KH_2_PO_4_ was used (5-125 mg/mL). Each isolate was tested in triplicate.

#### 2.5.3 Production of IAA

The ability of the NFB isolates to synthesize and secrete indol-3-acetic acid (IAA) was determined using the colorimetric method proposed by Ehmann (1977). The quantification protocol is described in Chen *et al*. (2017). Fresh cultures were adjusted to 0,5 McF, and 1 mL aliquots were added to nutrient broths supplemented with 200 mg/L of L-tryptophan (L-Trp). The tubes were incubated for five days at 30°C, after which the cell-free supernatant was recovered after centrifugation at 10000 *g* for 10 min. 1 mL of the supernatant was mixed with 0,1 mL of H_3_PO_4_ and 5 mL of Salkowski’s reagent. The mixture was kept in the dark for the next 30 min. The appearance of a pink color indicated the presence of IAA. The assay was repeated without L-tryptophan supplementation to test for the presence of Trp-independent biosynthesis pathways. Each isolate was tested in triplicate.

### 2.6 Pot experiments

#### 2.6.1 Experimental design

The study was divided into two experiments, one with bean plants (*P. vulgaris*), and another with rice plants (*O. sativa* var. Asp-18FL). Both experiments were carried out indoors, with daily temperatures ranging from 20 to 31 °C, and with a mean photoperiod of 12 h. Both experiments included the same six treatments: (i) seeds coated with BN03, (ii) seeds coated with BN09, (iii) seeds coated with BN16, (iv) seeds coated with BN18, (v) seeds coated with a consortium of the four PGPB, and (vi) seeds coated without inoculum. Two controls were included in the study: a positive control of seeds treated with 0,5% of Urfos-44 (Tripoliven C.A), which is a chemical fertilizer made up of phosphoric acid and urea (Pavone *et al*., 2020), and a negative control of seeds that were only irrigated with water. Each treatment and control included 25 plants, for which 10 *×* 15 cm plastic bags were used to grow the seed. The soil used was washed several times with sand to reduce its nutrient concentration, mimicking a low-fertility soil.

#### 2.6.2 Seeds and inoculum preparation

Bean and rice seeds were surface-sterilized and pre-germinated as described in 2.3.2. PGBP strains were grown in nutrient broth at 150 rpm until the exponential phase was reached (OD_600_ = 0, 6 − 0, 8). Bacterial cells were obtained by centrifugation at 7000 *g* for 10 min, and resuspended in 10 mM of MgSO_4_ · 7 H_2_O supplemented with 2% glycerol. Seed coating was achieved by mixing the bacterial isolates with 2% gum arabic in a 1:1 ratio, and slowly adding this mixture to the seeds. To complete bacterial adhesion, the seeds were air dried for 24 h at room temperature (Rocha *et al*., 2018).

#### 2.6.3 Sowing and harvest

Seeds were individually sown into their plastic bag. After 4 and 6 weeks of growth, the bean and rice plants were harvested, respectively. The shoot dry and fresh weight, the root dry and fresh weight, plant height, and number of leaves were determined. Dry weight was obtained by exposing plant matter to 60°C for 24 h. Germination was monitored during seven days on both types of plants, and the percentage and rate of germination (*R*) were determined using equations (2) and (3), according to Fiodor *et al*. (2023) and Maguire (1962), respectively:

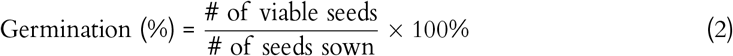

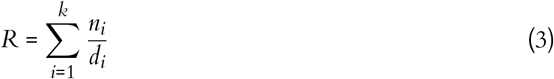

where *n*_*i*_ is the number of seeds that germinated on the *ith*-day, *d*_*i*_ is the *ith*-day after the experiment was started, and *k* is the number of days. Additionally, in the case of bean plants, the nodule biomass and symbiotic efficiency (SEF) were also determined. The latter was calculated as described in Gibson (1987). The SEF values were rated as: >80% = highly effective, 51–80% = effective, 35–50% = lowly effective and <35% = ineffective (Lalande *et al*., 1990).

### 2.7 Data analysis

Statistical analysis was performed with PAST (version 4.13). For the plant-growth promoting effects, a Kruskal-Wallis test was used in all cases to determine differences among the effects shown across isolates, with Dunn tests as post hoc tests. The same analysis strategy was used to examine the field experiment data. A significance level of α = 0, 05 was assumed in all tests.

## 3. RESULTS

### 3.1 Isolation and characterization of NFB isolates

Eighteen nitrogen-fixing bacterial strains were isolated from rhizosphere, nodule, and soil samples (Table I). 12 isolates are rhizobacteria (67%), 4 isolates correspond to rhizobia (22%) and 2 isolates are free-living bacteria (11%). The morphological and biochemical characterization of each isolate is also shown in Table I. 13 isolates correspond to Gram-negative bacilli (73%), 3 isolates correspond to Gram-negative coccobacilli, 1 strain is a Gram-positive diplobacilli (5,5%), and another strain is a Gram-positive streptobacilli (5,5%). None of the isolates were capable of undergoing cyst formation, and very few could produce spores (BN03, BN12, BN14, BN15, and BN17). Most of the isolates utilized sucrose and D-mannitol as carbon-sources. Other sugars were variably utilized. Most strains can also utilize organic salts (e.g. citrate), and some isolates metabolized all tested carbon sources, such as BN03, BN07, BN13, BN14, and BN15.

**Table I.**
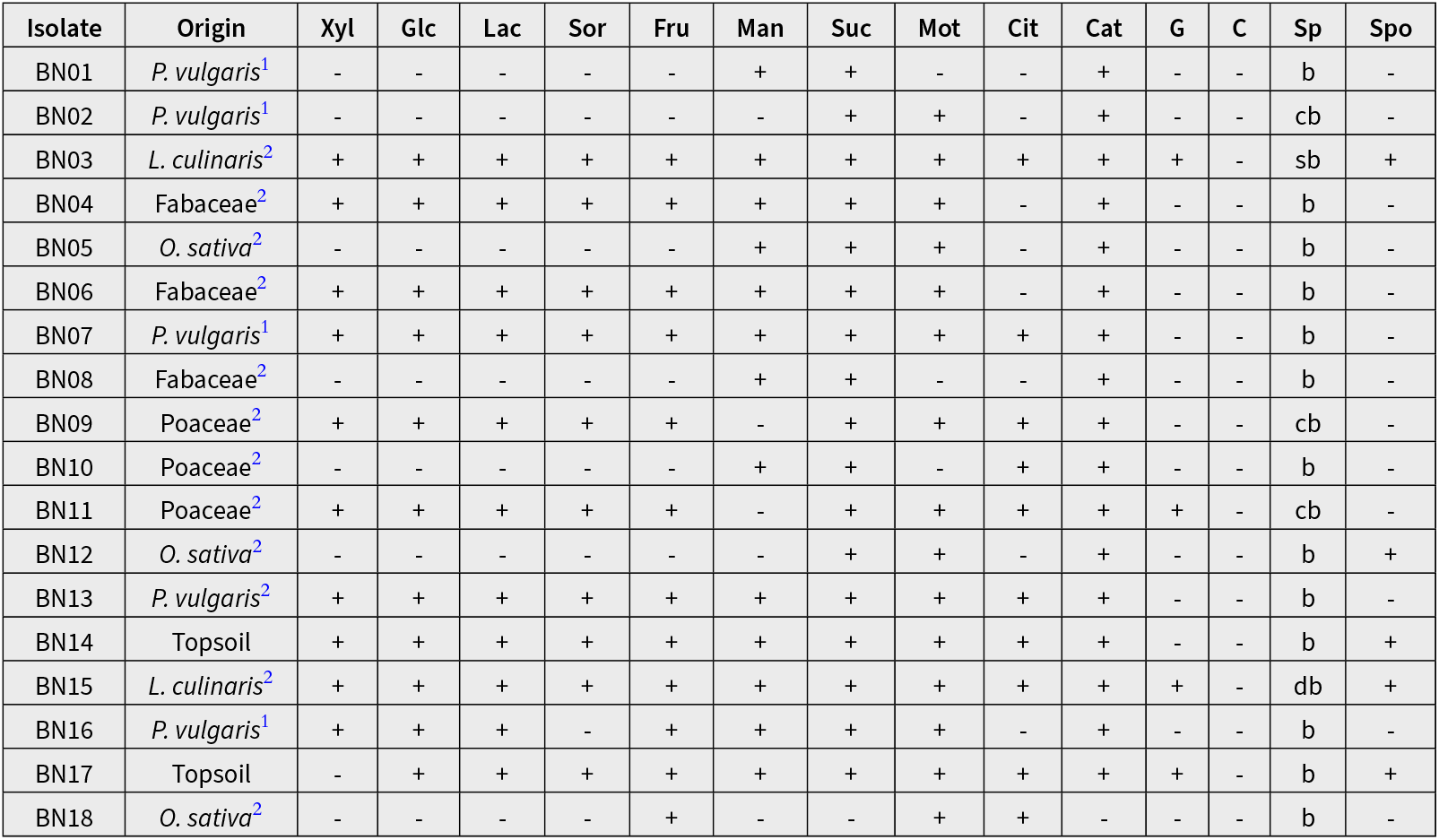
Biochemical profile of each isolate (Xyl = D-xylose, Glc = D-glucose, Lac = lactose, Sor = D-sorbitol, Fru = D-fructose, Man = D-mannitol, Suc = sucrose, Mot = motility, Cit = citrate, Cat = catalase, G = gram-type, C = cyst production, Sp = shape [b = bacillus, cb = cocobacillus, db = diplobacillus, sb = streptobacillus], and Spo = sporulation).^1^Isolated from nodules. ^2^Isolated from rhizopheres.

### 3.2. Screening for plant-growth promoting effects

Primary plant-growth promoting effects were assessed in every isolate (Table II). All strains exhibited nitrogen fixation, with most of them producing between 8-21 mg/L of ammoniacal nitrogen. Significant differences (*p* < 0, 0001) were found in fixation ability, with BN07 (19,43 mg/mL), BN09 (21,22 mg/mL), BN16 (25,14 mg/mL) and BN03 (27,88 mg/mL) exhibiting the highest fixation ability. The strain BN18 did not produce any NH_4_^+^ under aerobiosis, but showed luxuriant growth in NFb. In addition to nitrogen fixation, the ability of each strain to solubilize various forms of mineral nutrients was also assessed (Table II).

**Table II.** N_2_ -fixation and bacterial solubilization of P, K, and Zn. ^*a*^All results are expressed as mean ± sd. ^*b*^Corresponds to halo-assays. ^*c*^Calculated from Teles *et al.* (2024). ^*d*^Only fixes N_2_ in microaerobiosis.

| Isolate | N<sub>2</sub>-fixation<sup>a</sup> |  |  | Mineral solubilization<sup>b</sup> |  |  |  |  |
| --- | --- | --- | --- | --- | --- | --- | --- | --- |
|  | TAN (mg/mL) | PSI | Capacity | FSI | MSI | Capacity<sup>c</sup> | ZSI | Capacity |
| BN01 | 9,63 ± 2,90 | 0 | None | 3,43 ± 0,46 | 2,19 ± 0,18 | Medium | 0 | Low |
| BN02 | 10,49 ± 0,58 | 4,25 ± 0,94 | High | 0 | 0 | None | 0 | Low |
| BN03 | 27,88 ± 2,32 | 2,86 ± 0,12 | Medium | 2,15 ± 0,38 | 2,53 ± 0,31 | Medium | 1,81 ± 0,23 | Low |
| BN04 | 6,71 ± 2,97 | 3,22 ± 0,32 | Medium | 0 | 0 | None | 2,09 ± 0,35 | Medium |
| BN05 | 4,19 ± 1,63 | 0 | None | 0 | 0 | None | 0 | Low |
| BN06 | 15,16 ± 2,76 | 0 | None | 0 | 0 | None | 3,01 ± 0,89 | Medium |
| BN07 | 19,43 ± 2,72 | 5,30 ± 0,18 | High | 0 | 0 | None | 0 | Low |
| BN08 | 18,29 ± 1,75 | 4,04 ± 0,61 | High | 0 | 0 | None | 0 | Low |
| BN09 | 21,22 ± 1,06 | 2,88 ± 0,47 | Medium | 2,40 ± 0,38 | 4,27 ± 0,43 | High | 4,15 ± 0,18 | Medium |
| BN10 | 8,83 ± 2,25 | 0 | None | 0 | 0 | None | 0 | Low |
| BN11 | 10,02 ± 2,93 | 2,53 ± 0,30 | Medium | 0 | 0 | None | 0 | Low |
| BN12 | 8,90 ± 1,91 | 3,11 ± 0,21 | Medium | 0 | 1,27 ± 0,21 | Low | 0 | Low |
| BN13 | 13,71 ± 1,05 | 3,41 ± 0,44 | Medium | 0 | 0 | None | 0 | Low |
| BN14 | 6,20 ± 1,70 | 2,85 ± 0,56 | Medium | 0 | 0 | None | 2,13 ± 0,35 | Medium |
| BN15 | 4,17 ± 1,67 | 0 | None | 0 | 0 | None | 0 | Low |
| BN16 | 25,14 ± 1,87 | 4,56 ± 0,38 | High | 7,34 ± 0,91 | 3,64 ± 0,26 | High | 5,22 ± 1,59 | High |
| BN17 | 15 ± 1,55 | 2,80 ± 0,22 | Medium | 0 | 0 | None | 0 | Low |
| BN18<sup>d</sup> | — | 3,74 ± 0,39 | Medium | 5,41 ± 1,65 | 4,30 ± 0,33 | High | 2,03 ± 0,24 | Medium |

**Key:** TAN: total ammoniacal nitrogen; PSI: phosphorus solubilization index; FSI: feldspar solubilization index; MSI: muscovite solubilization index; ZSI: zinc solubilization index.

Thirteen nitrogen-fixing strains were not capable of solubilizing K-minerals (Table II). Feldspar was only solubilized by BN01, BN03, BN09, BN16, and BN18, who exhibited a medium to high solubilizing activity. The first three isolates only showed a medium capacity, but the latter two showed high capacity to solubilize the mineral. No difference was found between BN01, BN03 and BN09. The strain BN16 showed the highest potential, surpassing BN18 in capacity (*p* = 0, 016). BN12 was able to solubilize muscovite, but not feldspar, albeit with a low capacity.

Zn-solubilization was also not common among the nitrogen-fixing strains (Table II). Only BN03, BN04, BN06, BN09, BN14, BN16 and BN18 showed positive activity. Significant differences in solubilization ability were found between the isolates, with BN16 being the only one who exhibited high capacity. The rest of isolates mostly showed medium capacity, with the exception of BN03 showing low capacity (Table II).

Unlike K and Zn, P-solubilization was a more common trait among the nitrogen-fixing strains. More than two thirds of isolates were able to solubilize tricalcium phosphate. The strains BN01, BN05, BN06, BN10, and BN15 did not show any solubilization ability. The rest of the isolates exhibited medium to high activity, with the strains BN02, BN07, BN08, and BN16 showing the highest potential. Significant differences were also found between these, with BN07 and BN16 surpassing the other strains in solubilization capacity (*p* < 0, 05), with no differences found between the two. All isolates able to solubilize tricalcium phosphate also showed acidification of the surrounding media.

Besides the halo-based assay, the solubilization capacity of each isolate was quantitatively verified by the ascorbic acid method. The mean orthophosphate solubilized by the isolates is shown in Table III. Most isolates produced 45-90 µg/mL of phosphate. The strains BN01 and BN06 produced phosphate despite not showing any halo in the plate assay. 10 isolates showed a P-solubilization capacity above 70 µg/mL.

**Table III.** Production of orthophosphate and indol-3-acetic acid by the nitrogen-fixing isolates.

| Isolate | P-solubilization<sup>d</sup> | IAA production |  |
| --- | --- | --- | --- |
|  | PO<sub>4</sub><sup>-3</sup> (μg/mL) | L-Trp<sup>+</sup> | L-Trp<sup>-</sup> |
| BN01 | 20,66 ± 3,11 | + | - |
| BN02 | 74,73 ± 10,09 | + | + |
| BN03 | 80,60 ± 12,04 | + | - |
| BN04 | 72,93 ± 3,96 | + | - |
| BN05 | 0 | - | - |
| BN06 | 44,56 ± 8,05 | + | - |
| BN07 | 67,66 ± 14,06 | + | - |
| BN08 | 105,44 ± 4,75 | + | - |
| BN09 | 84,12 ± 6,87 | + | + |
| BN10 | 0 | - | - |
| BN11 | 41,78 ± 18,09 | + | - |
| BN12 | 70,25 ± 7,39 | + | + |
| BN13 | 93,64 ± 12,33 | + | - |
| BN14 | 64,67 ± 7,37 | - | - |
| BN15 | 0 | - | - |
| BN16 | 123,28 ± 6,97 | + | + |
| BN17 | 72,87 ± 3,33 | + | + |
| BN18 | 98,51 ± 3,28 | + | - |

Besides the production of orthophosphate, the capacity to produce and secrete indol-3-acetic acid with and without the supplementation of L-tryptophan was also evaluated (Table III). 15 nitrogen-fixing strains were capable of producing IAA when there was a supply of L-tryptophan. The biosynthesis of IAA without L-Trp supplementation was much rarer, as only five strains were able to produce the growth-regulator without the aminoacid, namely BN02, BN09, BN12, BN16, and BN17. Strains BN05, BN10, and BN15 did not produce IAA.

### 3.3 Selection of elite strains

Elite isolates were selected on the basis of their plant-growth promoting effects. A scoring system was used as the selection criteria (Table IV), where values were weighted and assigned to each promotion mechanism according to their contribution to plant-growth and development.

**Table IV.** Scoring system to identify the strains with the highest growth-promotion potential.

| Isolate | N<sub>2</sub>-fixation | P | K-fel | K-mus | Zn | IAA | Score |
| --- | --- | --- | --- | --- | --- | --- | --- |
| BN01 | + | - | ++ | ++ | - | + | 6 |
| BN02 | + | +++ | - | - | - | ++ | 7 |
| BN03 | + | ++ | +++ | ++ | + | + | 9 |
| BN04 | + | ++ | - | - | ++ | + | 6 |
| BN05 | + | - | - | - | - | - | 1 |
| BN06 | + | - | - | - | ++ | + | 4 |
| BN07 | + | +++ | - | - | - | + | 5 |
| BN08 | + | +++ | - | - | - | + | 5 |
| BN09 | + | ++ | ++ | +++ | +++ | ++ | 13 |
| BN10 | + | - | - | - | - | - | 1 |
| BN11 | + | ++ | - | - | - | + | 4 |
| BN12 | + | ++ | - | + | - | ++ | 6 |
| BN13 | + | ++ | - | - | - | + | 4 |
| BN14 | + | ++ | - | - | ++ | + | 6 |
| BN15 | + | - | - | - | - | - | 1 |
| BN16 | + | +++ | +++ | ++ | +++ | ++ | 14 |
| BN17 | + | + | - | - | - | ++ | 4 |
| BN18 | + | ++ | +++ | +++ | + | + | 11 |

Elite isolates were selected on the basis of their plant-growth promoting effects. A scoring system was used as the selection criteria, where values were weighted and assigned to each promotion mechanism according to their contribution to plant-growth and development. The system was developed with the purpose of identifying those strains with the greatest potential to function as a biofertilizer. The scoring system is as follows: one, two and three points were assigned to those strains with low, medium and high solubilization capacity, respectively. The ability to produce IAA with and without the supplementation of L-tryptophan was also awarded one point, and the same was done for the ability to fix atmospheric N_2_. The scoring system for the 18 strains is shown in Table IV. The isolates BN03, BN09, BN16, and BN18 scored the highest marks, and were thus selected for the field experiments.

### 3.4 Pot experiments

#### 3.4.1 Seedling phase

The germination rate and percentage of both bean and rice plants were determined after a week of development (Table V). The germination percentage of bean seedlings was higher in all bacterial treatments than the 65% observed in the negative control. An effect size of 5-10% was seen in the separate bacterial treatments, and up to 25% in the consortium. The combined action of the four strains proved to be more effective than any of the strains alone (*p* < 0, 05), with a germination percentage of 90%. No differences were found between the individual bacterial treatments (*p* > 0, 05).

**Table V.** Germination percentage and rate of bean and rice after seven days of growth. Values with the same letter do not differ significantly.

| Treatment | Germination (%) |  | Rate (seeds/day) |  |
| --- | --- | --- | --- | --- |
|  | Bean | Rice | Bean | Rice |
| BN03 | 70<sup>a</sup> | 75 | 9,25 | 5,83 |
| BN09 | 75<sup>a</sup> | 85 | 8,33 | 8,21 |
| BN16 | 75<sup>a</sup> | 85 | 9,17 | 6,75 |
| BN18 | 70<sup>a</sup> | 85 | 7,14 | 6,98 |
| Consortium | 90<sup>b</sup> | 70 | 10,45 | 6,68 |
| Gum arabic | 55<sup>a</sup> | 65 | 5,25 | 4,68 |
| Positive control | 80<sup>b</sup> | 85 | 8,5 | 6,12 |
| Negative control | 65<sup>a</sup> | 80 | 6,39 | 4,75 |

Germination rate can be used to evaluate the potential effect that bacterial strains have on early embryonic development. Each bacterial treatment achieved a faster germination speed compared to untreated bean seeds (Table V). The consortium exhibited the highest germination promotion, which was even higher than the germination rate of the seeds treated with Urfos-44. A different germination pattern was seen in the case of rice plants, as the consortium was also more effective than the individual bacterial treatments, which surpassed the consortium by 5-10%. Across all treatments, the bacterial strains accelerated the germination of rice seedlings (Table V). In either plant, the non-inoculated seeds coated with acacia gum showed the lowest germination percentage in the study (55-65%).

#### 3.4.2 Vegetative phase

The parameters measured after the harvest of bean plants are shown in Table VI. Each bacterial treatment significantly improved the shoot biomass, root biomass, plant height, and leaf number per plant when compared to the untreated plants of the negative control (*p* < 0, 05). The consortium showed to be the more efficient treatment in promoting the growth and development of the shoot system than any of the bacterial strains alone (*p* < 0, 05). However, this synergy was not expressed in the root development, as no differences were found between the consortium and the individual strains. None of the treatments affected plant hydration.

**Table VI.**
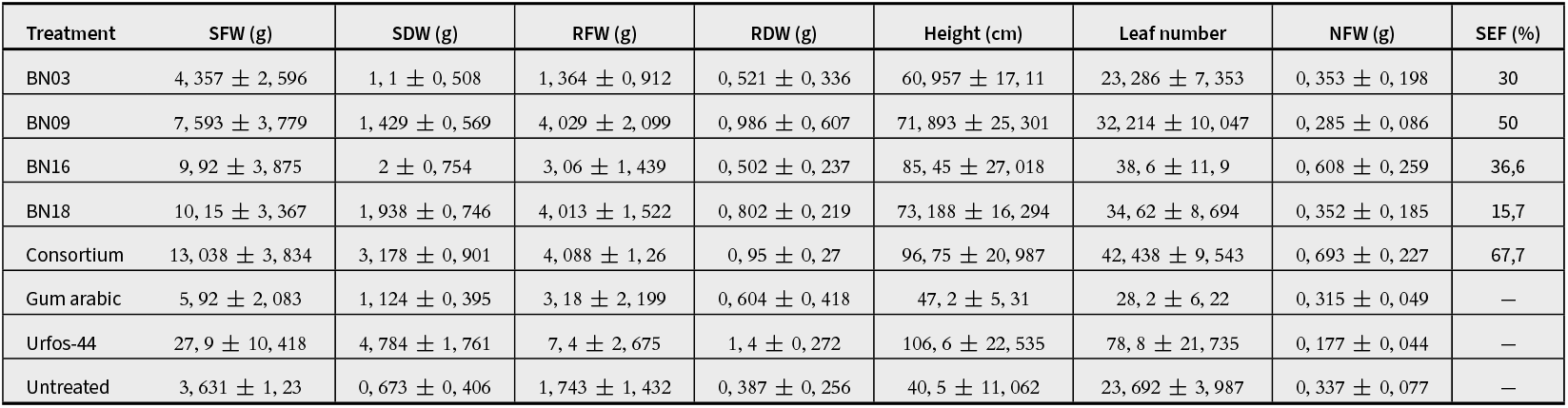
Effect of the bacterial treatments on the growth and development of bean plants. **Key:** SFW: shoot fresh weight; SDW: shoot dry weight; RFW: root fresh weight; RDW: root dry weight; NFW: nodule fresh weight.

| Treatment | SFW (g) | SDW (g) | RFW (g) | RDW (g) | Height (cm) | Leaf number | NFW (g) | SEF (%) |
| --- | --- | --- | --- | --- | --- | --- | --- | --- |
| BN03 | 4,357 ± 2,596 | 1,1 ± 0,508 | 1,364 ± 0,912 | 0,521 ± 0,336 | 60,957 ± 17,11 | 23,286 ± 7,353 | 0,353 ± 0,198 | 30 |
| BN09 | 7,593 ± 3,779 | 1,429 ± 0,569 | 4,029 ± 2,099 | 0,986 ± 0,607 | 71,893 ± 25,301 | 32,214 ± 10,047 | 0,285 ± 0,086 | 50 |
| BN16 | 9,92 ± 3,875 | 2 ± 0,754 | 3,06 ± 1,439 | 0,502 ± 0,237 | 85,45 ± 27,018 | 38,6 ± 11,9 | 0,608 ± 0,259 | 36,6 |
| BN18 | 10,15 ± 3,367 | 1,938 ± 0,746 | 4,013 ± 1,522 | 0,802 ± 0,219 | 73,188 ± 16,294 | 34,62 ± 8,694 | 0,352 ± 0,185 | 15,7 |
| Consortium | 13,038 ± 3,834 | 3,178 ± 0,901 | 4,088 ± 1,26 | 0,95 ± 0,27 | 96,75 ± 20,987 | 42,438 ± 9,543 | 0,693 ± 0,227 | 67,7 |
| Gum arabic | 5,92 ± 2,083 | 1,124 ± 0,395 | 3,18 ± 2,199 | 0,604 ± 0,418 | 47,2 ± 5,31 | 28,2 ± 6,22 | 0,315 ± 0,049 | — |
| Urfos-44 | 27,9 ± 10,418 | 4,784 ± 1,761 | 7,4 ± 2,675 | 1,4 ± 0,272 | 106,6 ± 22,535 | 78,8 ± 21,735 | 0,177 ± 0,044 | — |
| Untreated | 3,631 ± 1,23 | 0,673 ± 0,406 | 1,743 ± 1,432 | 0,387 ± 0,256 | 40,5 ± 11,062 | 23,692 ± 3,987 | 0,337 ± 0,077 | — |

The symbiotic efficiencies of each treatment are also presented in Table VI. The mean dry biomass of bean plants treated with BN16 and the consortium reached a minimum of 50% of the corresponding biomass of plants treated with the synthetic fertilizer. According to the classification of Lalande *et al*. (1990), the strain BN09 and the consortium used are then highly effective and effective biofertilizers, respectively. The remaining strains achieved lower symbiotic efficiencies, but they still significantly improved the growth of the bean plants treated.

When compared to the plants treated with Urfos-44, the bacterial treatments showed far less promotion is most biometric parameters, but not in all. The mean bean height in the consortium treatment was only slightly lower than mean height in the Urfos group. In most other metrics, the best performing bacterial treatments achieved 30-50% of the results of the Urfos-44 treated plants. The application of the acacia gum exclusively promoted the development of the roots of bean plants (Table VI), with no effect on any of the other plant metrics measured. The effect on the root system was also smaller than the one seen in most bacterial treatments (*p* < 0, 05), with the exception of treatment with BN03, where no differences were found.

Nodulation was not improved upon treatment with the BN03, BN09, and BN18 strains, as no differences were found when compared to the nodule biomass present in the untreated group (*p* > 0, 05), which represents the basal nodulation from the local rhizobia present in the soil used. In contrast, treatment with BN16 and the consortium significantly stimulated the nodulation in bean plants, with no difference between the two (*p* = 0, 531). Fertilization with Urfos-44 markedly reduced the number and size of nodules present in bean roots, with the lowest NFW of any of the treatments or controls.

Every bacterial treatment also improved every rice metric compared to the untreated plants of the negative control (*p* < 0, 05), although with a smaller effect size than that seen in bean plants (Table VII). In contrast to the bean assay: (i) no differences in shoot growth were found between the consortium and individual strains, BN16 and the consortium showed the highest shoot promotion, and (ii) the individual strains outperformed the consortium in root development, with strains BN09 and BN16 being the most effective.

**Table VII.** Effect of the bacterial treatments on the growth and development of rice plants. **Key:** SFW: shoot fresh weight; SDW: shoot dry weight; RFW: root fresh weight; RDW: root dry weight.

| Treatment | SFW (g) | SDW (g) | RFW (g) | RDW (g) | Height (cm) | Leaf number |
| --- | --- | --- | --- | --- | --- | --- |
| BN03 | 1,442 ± 0,75 | 0,279 ± 0,181 | 1,567 ± 1,088 | 0,306 ± 0,212 | 59,167 ± 5,149 | 6,5 ± 1,931 |
| BN09 | 2,055 ± 1,064 | 0,411 ± 0,212 | 2,318 ± 1,135 | 0,466 ± 0,228 | 61,956 ± 9,55 | 8,181 ± 2,136 |
| BN16 | 2,39 ± 0,686 | 0,48 ± 0,138 | 2,77 ± 0,715 | 0,549 ± 0,142 | 60,8 ± 5,827 | 9 ± 1,563 |
| BN18 | 1,151 ± 0,336 | 0,232 ± 0,081 | 1,36 ± 0,604 | 0,273 ± 0,121 | 57,3 ± 5,078 | 7,8 ± 1,619 |
| Consortium | 2,46 ± 0,785 | 0,504 ± 0,161 | 1,74 ± 0,412 | 0,358 ± 0,085 | 70 ± 9,775 | 10,5 ± 0,85 |
| Gum arabic | 1,12 ± 0,23 | 0,224 ± 0,046 | 1,18 ± 0,339 | 0,23 ± 0,066 | 52 ± 4,714 | 6,4 ± 0,843 |
| Urfos-44 | 9,243 ± 4,737 | 1,858 ± 0,952 | 4,014 ± 1,57 | 0,823 ± 0,322 | 81,714 ± 5,851 | 21,142 ± 5,368 |
| Untreated | 0,87 ± 0,325 | 0,174 ± 0,065 | 0,792 ± 0,296 | 0,159 ± 0,059 | 53,077 ± 4,212 | 6,308 ± 1,437 |

Treatment with BN16 and the consortium yielded a mean shoot biomass of 27% of the one achieved within the Urfos group. However, the gap between biofertilizers and Urfos-44 treated rice plants was smaller with respect to root development. For example, BN16 treatment achieved 68% of the root biomass of the synthetic group. The bacterial treatments also improved the height and number of leaves in the rice plants treated when compared to the control (*p* < 0, 05), with the consortium having had the biggest impact out of the treatments applied. The application of the acacia gum exclusively promoted the development of the rice roots, as was also observed in the bean group, without an effect on the other biometric parameters.

## 4. DISCUSSION

### 4.1 Characterization of isolates

The available carbon source is a key factor for microbial survival. Strains capable of utilizing various carbon sources are more likely to survive (Fiodor *et al*., 2023), especially in a new and often unfavorable environment, where the local microbiota is better adapted (Campos *et al*., 2014). In this study, most of the isolated nitrogen-fixing strains are capable of degrading multiple carbon sources, with some isolates being able to utilize all sources tested (e.g. BN03). This feature may boost their survival chances when used as a biofertilizer in field conditions (Martínez-Viveros *et al*., 2010). Motility can also be beneficial, especially under some delivery mechanisms. Most of isolated strains are motile, including the selected BN03, BN09, BN16 and BN18. Although these strains were immobilized to the seeds, their ability to move around could benefit the growing plant (Czaban *et al*., 2007).

The metabolic profile and morphological traits assessed allowed a preliminary identification of some of the isolated strains. Using *Bergey’s manual of systematic bacteriology* (Brenner *et al*. 2005), the strain BN03 likely belongs to the *Parabulkholderia* genus, a common nitrogen fixer of local soils (Artigas *et al*., 2021). The strain BN18 is with high confidence a member of *Azospirillum* genus, given that it only fixes N_2_ under microaerobiosis, and shows typical growth in RC agar (Cáceres, 1982). A member of the Rhizobiaceae family was also found among the isolated strains. The isolate BN16 is a rhizobia that could be assigned to the *Bradyrhizobium* genus, with typical growth in YMA, and the ability to nodulate *P. vulgaris* (Verástegui-Valdés *et al*., 2014). Strains BN09 and BN17 are likely members of the *Bacillus* genus, the latter strain showing great resemblance to *Bacillus subtilis*. The molecular identification of these isolates is recommended for future and similar studies.

### 4.2 Plant-growth promoting effects

Significant differences were found among the TAN values obtained from the isolates, which was expected given the differential growth observed during cultivation. Some isolates, namely BN03, BN07, BN08, BN09 and BN16, showed abundant growth in N-free media much faster than other strains, and these were the ones who produced the largest amount of ammoniacal nitrogen, between 18-28 mg/mL/week of TAN, which is higher than the TAN values reported in similar studies. For instance, Ha & Chu, (2020) isolated free-living diazotrophs that produced between 12-19 mg/mL of TAN, and a similar study by Nafisah *et al*. (2022) found rhizobacteria with TAN values of 8-15 mg/mL. The higher N_2_-fixation found in this study may be an indication of the potential of these strains to be used as biofertilizer.

The three major plant macronutrientes (N, P, and K) are largely considered to be limiting factors for the growth and development of cultivars, especially when considering current global food demands (Zhang *et al*., 2015; Walling & Vaneeckhaute, 2020; Yasmin *et al*., 2021; Fang *et al*., 2023), thus the ability of the nitrogen-fixing strains to solubilize and provide each of these macronutrientes was assessed. P-solubilization was common among the strains, with roughly 72% of the them being able to solubilize tricalcium phosphate (Table II). The ability of NFB strains to carry out mineral solubilization is well represented in the literature (Waday *et al*. 2021; Fiodor *et al*., 2023). Various mechanisms by which PGPB strains solubilize mineral nutrients have been proposed, which include organic acid production (e.g. citric acid, malic acid), chelation (e.g. siderophores, D-gluconic acid), enzyme secretion (e.g. phosphatases), and gas production (Glick, 2012). In this study, acidification of the surrounding media was seen in every case of P-solubilization, which is not unusual given that organic acid secretion has long been considered to be the typical mechanism of action (Nautiyal, 1999)

From the results obtained with the ascorbic acid method, it seems desirable that soil bacteria be screened with the NBRIP broth assay as opposed to the traditional halo-based techniques, as these assays are not reliable, given that in some cases strains who do not show any halo on agar plates can still solubilize less-readily forms of P (Nautiyal, 1999), as the strains BN01 and BN06 did. In contrast to phosphorus solubilization, K and Zn solubilization were not common activities among the isolates, with only a third or less exhibiting activity. The strain BN12 was only able to solubilize feldspar, but now muscovite, which may indicate nutrient-specific solubilization processes. Although organic acid production solubilized tricalcium phosphate, it was not sufficient to solubilize forms of K and Zn. The mechanisms of action by which the strains solubilized feldspar, muscovite and zinc oxide is unknown, and deserves further research.

Around 78% of strains were able to produce indol-3-acetic acid on the presence of its canon precursor L-tryptophan (Table III). The prevalence of auxin biosynthesis in PGPB has been well studied, and multiple lines of evidence indicate that around 80-90% of strains can secrete IAA through multiple metabolic pathways (Orozco-Mosqueda *et al*., 2023; Pérez-García *et al*., 2023), which suggests that this is one of the main mechanisms behind plant-growth promotion. Of the four selected strains, BN09 and BN16 can produce IAA with and without the presence of L-tryptophan, which is beneficial for plant-growth promotion because levels of L-tryptophan in soil are usually quite low.

### 4.3 Pot experiments

Improvements in germination rate were observed after treatment with the bacterial strains in both bean and rice seeds, compared to the untreated seeds of the negative control, which illustrated the potential of these PGPB strains to promote early embryonic development. The effect was not limited to the rate, as the percentage of germination also increased in the case of bean plants, by 5-10% in individual treatments and up to 25% in the consortium. Multiple studies have shown that these beneficial bacteria can accelerate germination (Poria *et al*., 2022; Pérez-García *et al*., 2023), allowing the rapid establishment of plant cultivars. This positive effect was seen in the bean group, but was limited to the germination rate in the rice group, which could be induced because rice is usually cultivated under water. PGPB largely impact germination by: (i) protecting the developing embryo through biocontrol mechanisms (Swarnalakshmi *et al*., 2020), and (ii) by the action of secreted growth regulators (e.g. IAA). The secretion of auxins has been considered as one of the main mechanisms by which these beneficial bacteria improve germination (Kour *et al*., 2019), as they stimulate the elongation and division of meristematic cells in the embryo (Wang & Ruan, 2013). As every selected strain produced IAA, even when L-tryptophan was not available (i.e., BN09 and BN16), the positive effect on germination may be due to this production.

The gum of various species have been used as an organic fertilizer in previous studies (Shobana *et al*., 2022) because of the positive net effect shown on the germination and growth of the plants (Shobana *et al*., 2022). However, the effect of acacia gum on plant development has not been studied. This gum is the standard seed-coating agent used to secure an effective delivery of beneficial microbes to the target plant (Zvinavashe *et al*., 2021). Although no significance was found when compared to the uncoated seeds, the application of acacia gum lowered the germination percentage by 10-15% in bean and rice plants. A possible explanation is that these complex polysaccharides may form a shell that limits seed imbibition, and thus extends dormancy. However, when paired with beneficial microbes, this effect may be compensated as germination is not handicapped, as observed in the NFB strains.

The bacterial treatments extended their effect beyond germination in both bean and rice, as improvements were observed in every biometric parameter when compared to the untreated plants. These improvements are due to the various plant-growth promoting factors that the selected strains posses. It is well known that N, P, K, and Zn are macronutrients that often limit the development and growth of cultivars due to their scarcity (Zhang *et al*., 2025; Fang *et al*., 2023), and whose deficiency worsens the productivity and quality of crops (Yasmin *et al* 2021). Millions of beneficial bacteria, like the strains isolated hereby, can provide these nutrients and promote growth as seen in this study and other studies (Waday *et al*. 2021; Fiodor *et al*., 2023), through atmospheric nitrogen fixation, nutrient mineral solubilization, growth-regulator production and secretion, among other mechanisms.

It is known that the use of chemical fertilizers can not be replaced by current biofertilizer and organic amendment technology because the production of these is not enough to sustain world growth demands (Liu *et al*., 2024). However, even a modest reduction in the production and use of chemical fertilizers would be beneficial (Moradzadeh *et al*., 2021; Mahmud *et al*., 2021), which together with other mitigation actions (e.g. production decarbonization) could significantly reduce their environmental and human-health impacts (Zhang *et al*., 2015). In the bean group, the mean biomass upon treatment with BN16 and the consortium achieved at least half of the corresponding biomass of plants treated with Urfos-44, without the use of any bacterial formulations (Campos *et al*., 2014). The effect was lower in the rice group, but treatment with BN16 still achieved 27% of the shoot biomass, and 69% of the root biomass of the Urfos group. The dual application of varying dosages of synthetic fertilizers together with biofertilizers under field conditions is recommended for future studies.

The application of acacia gum hampered the germination of both bean and rice seeds, but promoted their root development throughout their vegetative phase. The cause of this dual effect is not known. It may be that these polysaccharides limit seed imbibition while promoting rhizobacterial activity later on the development. A deeper analysis of its effect on plant development is needed, seeing how it is routinely used as the seed-coating agent in many biofertilizer applications.

Nodulation was improved in bean plants upon treatment with BN16 and the consortium, which was expected given that BN16 would correspond to a *Bradyrhizobium* strain (Favero *et al*., 2022). The lack of signficance between these two treatments suggests that the bulk of the effect is due to the nodulation induced by BN16. Nodulation was also significantly reduced by Urfos-44 application, as indicated by the number and size of the obtained nodules. This reduction has been reported in the literature, as previous studies have shown that excess N inhibits nodulation by reducing plant isoflavonoid production and the expression of *nod* factors within the rhizobia (Muzika, 1993; Barbulova *et al*., 2007), both factors being essential to the nodulation process. This interference with nodulation is one of the challenges of biofertilizer technology must face, particularly when using nitrogen-fixing strains. The use of formulations may minimize this inhibition.

The use of consortium is usually preferred to the use of single strains when formulating a biofertilizer (Zvinavashe *et al*., 2021), because multiple compatible strains can act synergistically to provide further nutrients and eliminate toxic products (Molina-Romero *et al*., 2017). In this study, the consortium only outperformed the single strains in some but not all parameters. However, it rarely showed worse performance than any of the individual treatments, thus it may be preferably used.

## 5. CONCLUSION

Eighteen nitrogen-fixing bacteria were isolated from rhizosphere, nodule and soil samples, all with the ability to fix atmospheric N_2_ in aerobiosis or microaerobiosis. 13 nitrogen-fixing isolates were found to solubilize tricalcium phosphate, 5 strains were found to solubilize less-readily available forms of K, and 6 strains solubilized zinc oxide. P-solubilization was likely carried out through the acidification of the surrounding media by organic acid secretion. However, such acid production was not able to solubilize feldspar, muscovite or zinc oxide. The near totality of nitrogen-fixing strains were able to produce IAA under the presence of L-trytophan, and 5 strains were able to also biosynthesize it in the absence of the precursor. The production of IAA by Trp-independent pathways is a desirable trait in PGPB formulation, as levels of L-trytophan in soil are tipically low.

The isolates BN03, BN09, BN16 and BN18 were selected on the basis of their plant-growth promoting traits, and were used in pot experiments with bean and rice plants. No formulation was used and seed-coating was used as the delivery mechanism. These four strains were applied separately and as a consortium in both groups. All five bacterial treatments significantly improved the shoot biomass, root biomass, plant height, and number of leaves per plant when compared to the untreated seeds. The effect was larger in bean plants, likely due to the nodulation induced by BN16, a putatative *Bradyrhizobium* strain. The treatment with BN16 and the consortium achieved 67% and 50% of the biomass of plants evaluated with Urfos-44, which indicates the potential that these strains have to promote the growth and development of important crops. The treatment with BN16 yielded 27% of the shoot biomass, and 68% of the root biomass of plants treated with Urfos-44. The application of acacia gum induced a dual effect on germination and plant development. Although root development was favored in both bean and rice plants upon treatment, germination rate and percentage was lowered in all cases. A possible explanation is that these polysaccharides form a shell on the seeds during germination which reduces seed imbibition, and thus extends dormancy, but they promote rhizobacterial activity during later growth stages, which promotes root development. Further research on the effects of acacia gum on plant development is needed.

These results, along with other previous work, indicate the potential that PGPB technology has to mitigate the environmental and human-health risks that the production and use of chemical fertilizers cause, promoting the transition towards sustainable global food production, where agricultural demands are met while protecting our environment and health.

